# Posttranscriptional activity of the eukaryotic translation initiation factor eIF4E contributes to HoxA9-driven leukemogenesis

**DOI:** 10.1101/2025.02.10.637540

**Authors:** Fang Zhou, Biljana Culjkovic-Kraljacic, Christian Bach, Li Feng, Yuta Mishima, Katherine L. B. Borden, Daniel G Tenen

**Author notes:** <u>co-last/co-corresponding authors</u>, **Correspondence** Daniel G. Tenen, Harvard Stem Cell Institute, Center for Life Science Room 437, Boston, MA, USA., Katherine L Borden, Department of Pharmacology and Robert H. Lurie Cancer Centre, Northwestern University, Chicago, Illinois, USA. <u>co-first authors</u> (Fang Zhou, Biljana Culjkovic-Kraljacic).

## Abstract

HoxA9, a homeodomain-containing transcription factor, is mis-expressed in over half of acute myeloid leukemia (AML) cases, and is associated with poor prognosis. Previous studies indicated that HoxA9 binds to the eukaryotic translation initiation factor eIF4E in primary specimens and that HoxA9 stimulated the RNA export and translation efficiency of selected RNAs via eIF4E. However, the relevance of this to its leukemogenic transformation capacity was unknown. Here, we used a double point mutation (HoxA9AA) to disrupt the physical and functional interaction between eIF4E and HoxA9 while retaining the HoxA9 transcriptional signature. Surprisingly, the mutation dramatically increased AML latency from a median of 90 to 280 days and resulted in incomplete penetrance. Re-transplantation of bone marrow cells from leukemic animals demonstrated even more pronounced differences in disease kinetics and penetrance with all animals succumbing to disease by day 60 in the wildtype group, while some HoxA9AA mice never developed leukemia. Collectively, these findings uncover a novel, transcription-independent mechanism of HoxA9-driven leukemogenesis through eIF4E and positions eIF4E as a potential therapeutic target AML patients expressing high levels of HoxA9.

**Key Points:**

- A double point mutation in HoxA9 disrupted the physical and functional interaction between eIF4E and HoxA9 while retaining the HoxA9 transcriptional signature.
- Eukaryotic translation initiation factor eIF4E contributes to HoxA9-driven leukemogenesis and is important for the maintenance of acute myeloid leukemia.

## Introduction

Homeobox A9 (HoxA9) is mis-expressed in ∼70% of AML cases and associated with poor outcomes^1,2^. HoxA9 modulates cell stemness and differentiation in normal hematopoiesis, showing a substantial transition from high expression in hematopoietic stem cells to undetectable in mature differentiated cells^3,4^. These activities are attributed to its well-established role as a homeobox transcription factor whereby it binds DNA. However, HoxA9 participates in activities beyond transcription, and their contribution to its leukemogenic capacity is unknown. An understanding of this is important for future development of therapies directed at HoxA9. HoxA9 physically and functionally interacts with the eukaryotic translation initiation factor eIF4E^5,6^. eIF4E is dysregulated in a subset of AML patients in which it is elevated, and enriched in the nucleus^7-10^. This corresponds in patients to inappropriate elevation of eIF4E-depedent nuclear mRNA export of oncogenic mRNAs^7-10^. Indeed, there is a strong link between eIF4E-dependent nuclear RNA export and its oncogenic capacity^8-14^. eIF4E has been successfully targeted in early-stage AML clinical trials with the antiviral drug ribavirin, which acts as a competitor of its normal ligand, the m^7^G cap on RNAs^8-10^. Ribavirin targeting of eIF4E corresponds to objective clinical responses, including complete remissions in a subset of high-eIF4E AML patients^8-10^. Interestingly, eIF4E and HoxA9 physically interact in both biophysical studies and in primary high-eIF4E AML patient specimens, and HoxA9 stimulated eIF4E-dependent nuclear RNA export and translation of oncogenic RNAs^15^. However, the biological impact of these interactions have not been elucidated. Here, we demonstrate for the first time that HoxA9 relies, at least in part, on eIF4E for its leukemogenic capacity, independent of the role of HoxA9 in transcription.

## Methods

### HoxA9 and eIF4E biochemistry assays

The HoxA9AA mutant was generated using QuikChange site-directed mutagenesis (Agilent, La Jolla, CA) and validated by sequencing. GST pulldowns, RNA export assays and western blots were as described^12^.

### Animal models

Retroviral supernatants for MIG-flag-HoxA9, MIG-flag-HoxA9AA, and MIY-Meis1 vectors were prepared as described^16^. 20,000 lin-/Sca1+/c-Kit+(LSK) cells/recipient were harvested from C57B/6 mice (CD45.2+) and transduced with a combination of Meis and HoxA9wt or HoxA9AA retroviral supernatants as described^16^. Transduced LSK were mixed with 200,000 normal bone marrow cells from B6.SJL-Ptprc^a^Pepc^b^/BoyJ (CD45.1+) mice per recipient and injected intravenously into lethally irradiated (9 Gy) B6.SJL-Ptprc^a^Pepc^b^/BoyJ mice. Cells were analyzed by flow cytometry. Serial replating assays were as in^17^.

### Microarray

BMCs from 3 Hoxa9wt/Meis1and 3 Hoxa9AA/Meis1 mice were harvested with ≥ 97% GFP+ cells in the leukemic bone marrow. The Mouse Gene 1.0 ST arrays were processed using the ‘oligo’ Bioconductor package, quality-controlled with arrayQualityMetrics, and RMA normalized. Data are deposited (GEO GSE288783).

### Statistics

Data were analyzed with the IBM SPSS statistics 26 and GraphPad Prism 8 software.

## Results and Discussion

To probe the relevance of eIF4E to the leukemogenic potential of HoxA9 *in vivo*, we generated a HoxA9 mutant targeting its eIF4E-binding motif, changing residues 11 and 16 in the wild type (11-**Y**VDSF**L**L-17) to alanine (11-**A**VDSF**A**L-17, HoxA9AA) (Figure 1A). The purified mutant GST-HoxA9AA bound to endogenous eIF4E with a ∼2-fold reduction relative to wild-type HoxA9 (HoxA9wt) in K562 cells. No interaction was observed with the negative control, eIF4E binding-protein1 (4E-BP1) (Figure 1B). GST-eIF4E pulldowns were also conducted using K562 cell lines stably expressing HoxA9wt or HoxA9AA. The mutation impaired eIF4E-binding by 2-fold relative to HoxA9wt, with no binding to the vector or GST controls. In this case, 4E-BP1 served as a positive control binding to eIF4E, and did not bind GST, as expected (Figure 1C). GST protein loading was equivalent for all experiments (Supplemental Figure 1A, Supplemental Figure 1B). This mutant and wildtype HoxA9 protein bound to DNA as shown in electrophoretic mobility assays, indicating that the mutation did not disrupt its DNA binding capacity (Supplemental Figure 1C).

**Figure 1.**
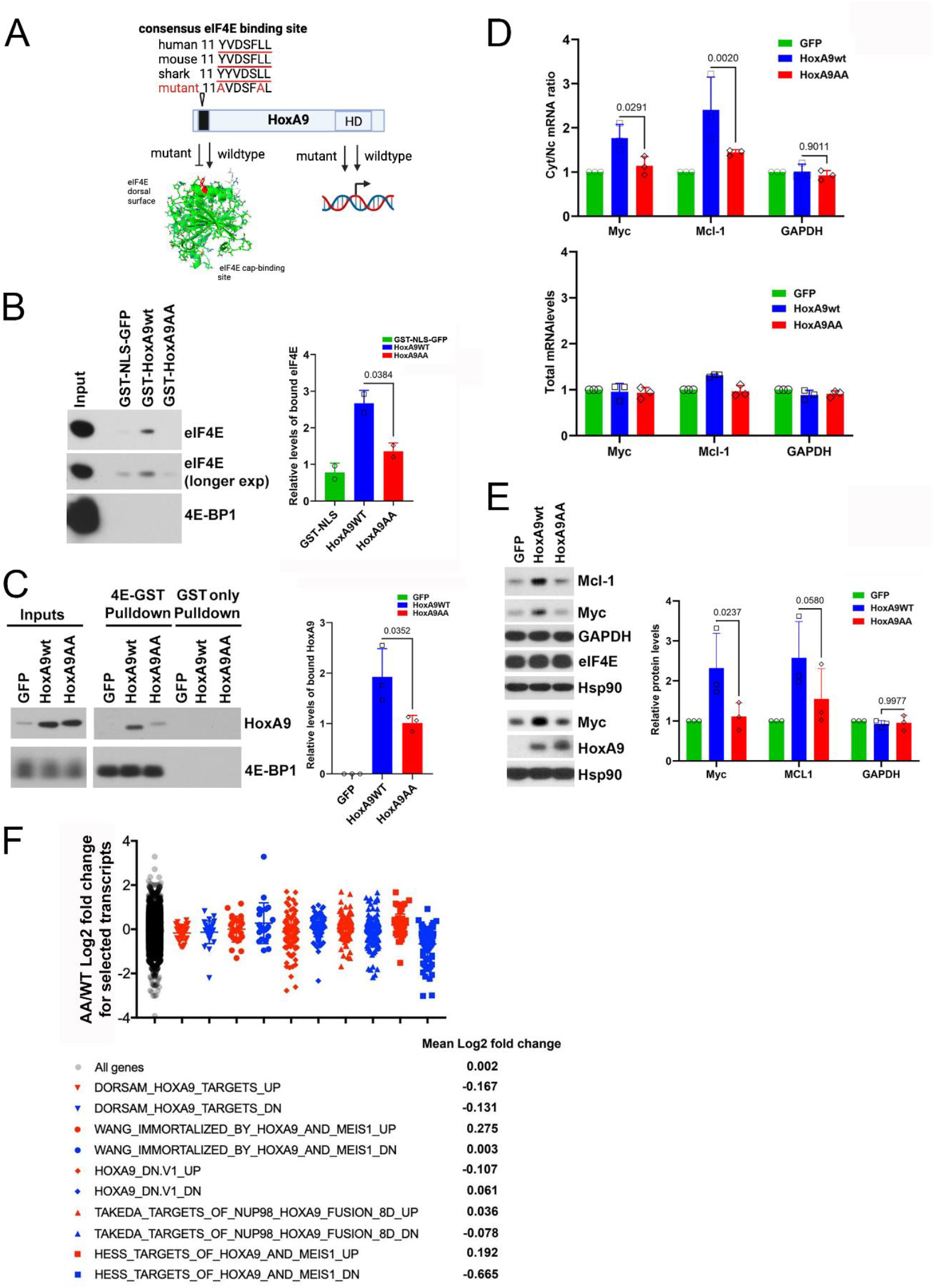
Mutation in the HoxA9 eIF4E-binding site reduces eIF4E interaction without affecting transcriptional functions of HoxA9 (A) Schematic depiction of the HoxA9 protein, showing consensus eIF4E binding site (black bar) that is conserved from humans to sharks. This site is defined by 11 YVDSFLL 17 and F is any hydrophobic residue. The mutant is generated by alanine substitutions of the conserved Y and L within this site and denoted HoxA9AA to disrupt eIF4E binding. Using purified proteins, HoxA9 directly binds the dorsal surface of eIF4E with a point mutation of W73 impairing binding^15^ (shown in red). Structure image made in pymol, pdb 2GPQ (apo-form of eIF4E). HD indicates homeodomain. Both mutant and wildtype bind DNA (Supplemental Figure 1C). (B) Immunoblot to assess the binding of purified GST-HoxA9 to eIF4E from K562 cell lysate by GST pulldown. Right panel is quantification over 3 independent replicates. Standard deviations and the means are shown. (C) Immunoblot to access the binding of HoxA9wt and HoxA9AA from K562 lysates to purified GST-eIF4E by GST pulldown. Right panel, quantification of three replicates. Means are shown and error bars indicate standard deviation. (D) HoxA9 promotes eIF4E-dependent mRNA export. Top panel: relative cytoplasmic/nuclear ratio of *MYC* and *MCL1* mRNAs in HoxA9 overexpressing cell lines. Fractionation controls are shown in Supplemental Figure 1D. Bottom panel: total *MYC* and *MCL1* mRNA levels in HoxA9 overexpressing cells. (E) Immunoblot to assess the protein expression of eIF4E-dependent RNA export targets in HoxA9 overexpressing cells. Right panel: quantifications over three replicates. Means are shown and error bars indicate standard deviation. P-values are provided. (F) Analysis of fold change in gene expression of HoxA9AA/Meis1 versus HoxA9wt/Meis1-induced leukemia demonstrated very few changes consistent with these constructs having similar transcriptional activity as observed from microarray analysis. Leukemic cells were obtained from 3 different mice per group. Ratio of HoxA9AA/HoxA9wt is shown. An analysis of all genes from comparison of the microarray is provided. The expression levels of HoxA9-regulated gene sets, selected from GSEA database, were analyzed for overlapping genes in our microarray dataset, and mean Log2 fold change was determined for each gene set.

To examine the functional impact of this mutation, we first monitored the capacity of HoxA9 to stimulate eIF4E-dependent nuclear mRNA export and production of MYC and MCL1 (Figure 1D)^13,14,18^. Cells were fractionated into nuclear and cytoplasmic compartments, and RNA content measured using RT-qPCR. HoxA9wt stimulated mRNA export of *MCL1* by nearly 3-fold and *MYC* nearly 2-fold relative to vector controls (Figure 1D). The HoxA9AA mutation reduced export by ∼2-fold for *MCL1* and ∼1.6-fold for *MYC* versus HoxA9wt. Consistently, we observed that HoxA9wt stimulated production for both MYC and MCL1 proteins by > 2-fold while Hox9AA was similar to vector controls as observed by western blot (Figure 1E). Total RNA levels were unchanged for *MCL1, MYC*, or *GAPDH* indicating these are not transcriptional effects of HoxA9 (Figure 1D, lower panel) and fractionation controls are provided (Supplemental Figure 1D). GAPDH protein levels and RNA export were not affected by either HoxA9wt or HoxA9AA. Thus, reduced stimulation of eIF4E activity by HoxA9 (∼2-3 fold decreased mRNA export and protein production of eIF4E targets for mutant) is similar to the decrease in the eIF4E-HoxA9 interaction (∼2 fold lower binding for mutant).

To assess whether the HoxA9-eIF4E interaction affected biological effects of HoxA9 *ex vivo*, we monitored plating efficiency of sorted primary murine HSPCs (LSK cells, Lin-/Sca-1+, c-Kit+) by performing serial re-plating assays in cytokine supplemented methylcellulose after retroviral transduction of freshly isolated murine HSPCs with HoxA9wt or HoxA9AA with the cofactor Meis1. After 4 rounds of re-plating, the colony forming potential of HoxA9AA-transduced cells was reduced by ∼2.8-fold relative to HoxA9wt (Figure 2A).

**Figure 2.**
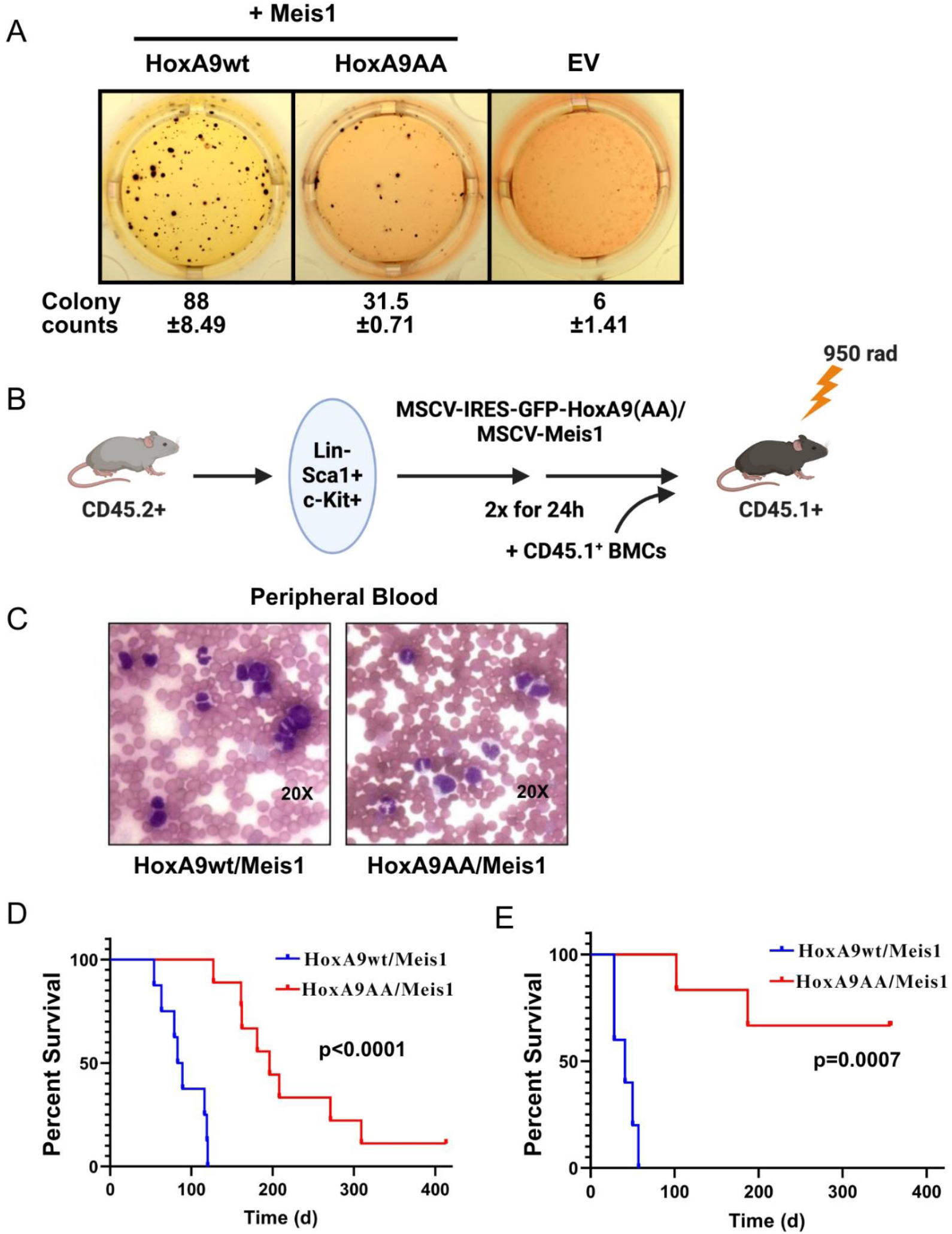
Reduction of the HoxA9-eIF4E interaction impairs leukemogenesis. (A) Colonies of primary lin-Sca-1+c-Kit+ (LSK) cells transduced with HoxA9/Meis1, HoxA9AA/Meis1 and corresponding vector controls after 4 rounds of serial replating in cytokine supplemented semi-solid medium. The mean number of colonies and the standard deviation are shown below. EV indicates empty vector. (B) Schematic presentation of murine transplantation model. (C) Peripheral blood smears (Wright-Giemsa stained) with magnified sections exemplifying colony morphology. Magnification: 20 × 1 . (D) Kaplan-Meyer plot of mice tranplanted with HoxA9/Meis1 (n=10) or HoxA9AA/Meis1 (n=14) transduced LSK cells. (E) Kaplan-Meyer plot after secondary transplants of HoxA9wt/Meis1 or HoxA9AA/Meis1 leukemic bone marrow cells (see Supplemental Figure 2) into secondary recipient mice (n = 5 per group).

We next utilized this mutant to evaluate the contribution of eIF4E to HoxA9-induced leukemic transformation *in vivo*. We transplanted CD45.2+ LSK cells transduced with Meis1 and either HoxA9wt-GFP or HoxA9AA-GFP into lethally irradiated CD45.1+ recipient mice (Figure 2B). Transduction efficiency was similar, 22.9% and 24.5%, for HoxA9wt/Meis1 and HoxA9AA/Meis1, respectively (Supplemental Figure 2A). There was comparable engraftment with similar kinetics in HoxA9wt/Meis1 and HoxA9AA/Meis1 transduced cells as seen by similar fractions of CD45.2 positive and GFP positive peripheral blood cells at four weeks post transplantation (Supplemental Figure 2B). Both groups eventually developed myeloid leukemia with 97-99% CD45.2+/GFP+ in the bone marrow of affected mice (Supplemental Figure 2C). Histologically, peripheral blood smears of affected mice revealed an extensive myeloblastic population with both mature and immature blast cells in the periphery consistent with myeloid leukemia and observed markers for the myeloid lineage, but not for B- or T-lymphoid lineages (Figure 2C, Supplemental Figure 2C). Despite the phenotypic similarities between HoxA9wt and HoxA9AA induced leukemias, there were highly significant differences in latency of disease onset and penetrance. All animals (n=10) in the HoxA9wt group succumbed to aggressive leukemia within 120 days (median 90 days), corresponding to full penetrance of HoxA9wt/Meis1 leukemia (Figure 2D). By contrast, the median latency in the HoxA9AA/Meis1 group (n=14) was 280 days. Moreover, two HoxA9AA/Meis1 mice never developed leukemia. One of these mice had detectable but low frequency of GFP positive cells in peripheral blood and bone marrow when sacrificed 413 days post transplantation. The second mouse had detectable GFP-positive peripheral blood cells until day 175 post-transplantation, but no remaining GFP+ cells in the bone marrow at sacrifice on day 190.

We next monitored the impact of leukemic cells in secondary recipients. Re-transplantation of bone marrow cells from leukemic animals demonstrated an even more pronounced difference in latency and penetrance in secondary recipients (Figure 2E). While 5/5 HoxA9wt/Meis1 transplanted secondary recipients developed myeloid leukemia and died within 60 days after transplantation, only 2 of 5 in the HoxA9AA/Meis1 group developed disease even after a year post-transplantation. The remaining 3 animals had no detectable GFP+ cells in peripheral blood or bone marrow. This strongly indicates that the eIF4E interaction is continuously required for the maintenance of efficient leukemic transformation by HoxA9. Thus, eIF4E is critical for HoxA9 mediated induction and maintenance of leukemogenesis. Furthermore, eIF4E-dependence is not regularly overcome by leukemic evolution, highlighting eIF4E inhibition as a potential target for therapeutic intervention in high HoxA9 AML.

Central to the dogma of HoxA9 functionality is its role in transcription. To investigate if HoxA9AA induced a different transcriptional program to HoxA9wt, we analyzed global mRNA expression profiles in bone marrow using specimens from leukemic wildtype and mutant mice. Microarray analysis revealed no major differences in the HoxA9 gene expression signatures between HoxA9wt and HoxA9AA transduced leukemia cells using MSigDB^**19**^ (Figure 1F). Indeed, of the 23,589 transcripts observed, only 1.7% of transcripts changed with 163 transcripts significantly downregulated, and 247 upregulated in HoxA9AA versus HoxA9wt, respectively (>1.5 fold change, p-value<0.05). These were not enriched in specific ontology pathways. These data strongly suggest that the reduced leukemogenic capacity of HoxA9AA is not caused by modified transcriptional activity. Consistent with these observations, the HoxA9AA mutant demonstrated DNA binding activity similar to that of wildtype (Supplemental Figure 1C).

In conclusion, our study illustrates a new mechanism underpinning HoxA9-mediated leukemogenesis. We demonstrate that even though HoxA9AA elicits a highly similar transcriptional program as HoxA9wt and binds DNA, this mutation substantially extends disease latency and reduces disease penetrance. Indeed, the mutation highlights that the HoxA9 interaction with eIF4E plays an essential role in induction and maintenance of HoxA9-dependent leukemogenesis. Strategies to directly target HoxA9 in patients remain elusive. Our findings provide an interesting alternative whereby AML patients with dysregulated HoxA9 could benefit from targeting eIF4E with drugs like ribavirin.

## Supporting information

Supplemental Methods, References, and Figures

## Acknowledgements

We are grateful for assistance from Dr Laurent Volpon for purified proteins. This work was supported by the NIH (RO1 98571 and RO1 80728 to KLB; 1P01HL131477-6 A1 to DGT); the Canada Research Chair programme (KLB); and the AGA/Jenzabar Foundation (DGT).

## Authorship contributions

FZ, BC-K, KLB, and DGT designed the experiments; CB, LF, and YM helped with design and performance of the experiments; FZ, BC-K, KLB, and DGT wrote the manuscript.

