## Supplemental Methods, References, and Figures for "Posttranscriptional activity of the eukaryotic translation initiation factor eIF4E contributes to HoxA9-driven leukemogenesis"

### Supplemental Methods and Materials

#### 1. Cell lines

K562 (CCL-243) were maintained in RPMI-1644 medium (Gibco) supplemented with 10% FBS. All cells were incubated at 37°C in a humidified atmosphere of 5% $CO_2$ on standard tissue culture plates and were routinely checked to ensure that there was no mycoplasma contamination using MycoAlert Mycoplasma Detection kit (Lonza,NY, U.S.A, LT07-418). Cell lines were authenticated by STR profiling (Wyndham Forensic Group). Primary murine HoxA9/Meis1 and HoxA9AA/Meis1 leukemia cell lines were established by harvesting BMCs from leukemic mice. 100,000 isolated and washed cells were subsequently cultured in RPMI+10% FBS (HyClone) supplemented with 100 ng/ml murine recombinant SCF and 10 ng/ml of each murine recombinant IL-3, IL-6, and GM-CSF (StemCell Technologies). After 1 week in culture, the cytokine supplementation was reduced to 10 ng/ml SCF and 5 ng/ml of each IL-3, IL-6, and GM-CSF and subsequently (after another week) to 2.5 ng/ml of each SCF, IL-6, and GM-CSF and 5 ng/ml IL-3. Finally, after another week cells were cultured in RPMI+10% FBS with 5 ng/ml murine recombinant IL-3 and used for experiments.

### 2. Wright-Giemsa staining

Cell morphology was evaluated by Wright-Giemsa staining after cytospin preparation of BMCs (1,000 rpm for 60s) or peripheral blood smear. Modified Wright-Giemsa Staining (Siemens Healthcare Diagnostics) was carried out according to the manufacturer's instructions. Image acquisition was carried out on an Axioscope 2 microscope (Zeiss).

### 3. Western blotting

Whole-cell extracts were lysed in RIPA lysis buffer (50 mM Tris-HCl pH 7.5,150 mM NaCl, 0.1% SDS, 0.5% sodium deoxycholate, 1% Triton X-10 1% Triton X-100, 5mM EDTA, 1X protease inhibitor cocktail) as described[1]. Lysates were then spun at 12000 rpm for 10 min to remove insoluble material and 20 µg of protein samples were loaded and resolved on a 10% SDS polyacrylamide gels. After electrophoresis, the proteins were transferred to a PVDF membrane using Trans-Blot® Turbo™ Transfer System (Bio-Rad Laboratories), blocked with 5% milk in TBS with 0.1% Tween-20, and probed with antibodies of interest overnight at 4°C. The blots were washed with TBS with 0.1% Tween-20, incubated with HRP-conjugated secondary antibodies for 1h, and finally examined by chemiluminescence using Supersignal west pico plus HRP substrate (ThermoScientific). Antibodies used for immunoblotting: mouse monoclonal antieIF4E (BD PharMingen), mouse monoclonal anti-β-actin (A5441 Sigma Aldrich), rabbit polyclonal anti-Mcl-1 (S19, sc819 Santa Cruz), rabbit polyclonal anti-Myc (ab32072 Abcam), mouse monoclonal anti-Hsp90 (sc-69703 Santa Cruz), rabbit polyclonal anti-GAPDH (sc-25778 Santa Cruz), rabbit monoclonal 4E-BP1 (53H11 Cell Signaling), and rabbit polyclonal anti-HoxA9 (NBP2-24633 Novus).

**4. GST pulldown**

GST-eIF4E, GST-NLS-GFP, GST only, GST-HoxA9wt and the GST-HoxA9AA mutant were purified as previously described [2,3]. Purified proteins (20 μg) were bound to 25μL of pre-equilibrated Glutathione Sepharose 4B beads (GE Healthcare) for 1h at 4°C with rotation in binding buffer (10 mM sodium phosphate pH 7.5, 50 mM NaCl, 1 mM DTT, 1.5mM $MgCl_2$ and 0.02% Nonident-P40, 1X protease inhibitor cocktail), and washed 3 times in the same buffer. K562 cells overexpressing GFP and HoxA9wt or HoxA9AA proteins were lysed in binding buffer containing protease inhibitors. Cell lysates were pre-cleared with 25μl pre-equilibrated (in Binding buffer) Glutathione Sepharose 4B beads (GE Healthcare) for 30 min at 4°C with rotation. 250μg of cell lysates were mixed with corresponding GST bound proteins in 500ml final volume and incubated for 1h at 4°C with rotation. Unbound protein was removed by washing six times with binding buffer, and bound proteins were eluted with Laemmli sample buffer for 5min at 95°C. The eluted protein complexes were then separated by SDS-PAGE and revealed using specific antibodies and Western blot analysis. Loading for purified proteins and inputs was monitored by Ponceau S staining. All experiments were carried out at least three independent times.

GST-NLS-GFP, GST-HoxA9wt and GST-HoxA9AA proteins were incubated with lysates from GFP transduced K562 cells and resulting pull-downs were probed with eIF4E antibody. GST-eIF4E or GST only were incubated with lysates from GFP, HoxA9wt or HoxA9AA overexpressing K562 cells and probed with HaxA9 antibody.

### 5. Cellular fractionation and RNA export assay

Cellular fractionation was carried out as previously described[1]. Briefly, ~$3\times10^7$cells were collected and washed twice in ice cold PBS (1200 rpm, 5min) and then re-suspended with slow pipetting in 1ml of Lysis buffer B (10 mM Tris pH 8.4, 140mM NaCl, 1.5mM $MgCl_2$, 0.5% NP40, 1mM DTT and 100 U/ml RNaseOUT (ThermoFisher Scientific). The lysates were centrifuged at 1000 g for 3 min at 4°C and supernatant (cytoplasmic fraction) was transferred into a fresh microtube. The pellet (nuclear fraction) was resuspended in 1 ml of Lysis buffer B, transferred to round bottom polypropylene tube. Then, 1/10 volume (100µl) of detergent stock (3.3% (w/v) Sodium Deoxycholate, 6.6% (v/v) Tween 40 (in DEPC $H_20$) was added with slow vortexing, and incubated on ice for 5 min, transferred to a microtube and centrifuged at 1000 g for 3 min at 4°C. Supernatant(post-nuclear fraction) was transferred to a fresh tube and the pellet-nuclear fraction was rinsed in 1 ml of lysis buffer B and centrifuged at 1000 g for 3 min at 4°C. The postnuclear and cytoplasmic fractions were combined. The RNA was extracted from the different fractions by adding TRIzol reagent (ThermoFisher Scientific) and Direct-zol RNA Miniprep Kit (Zymo Research) according to the manufacturer's instructions. The quality of fractionation was assessed by semi-qRT-PCR using U6snRNA and $tRNA_{Met}$ primers for nuclear and cytoplasmic fractions, respectively.

**6. RNA extraction and quantitative PCR**

DNAse treated RNA samples (Direct-zol RNA Miniprep Kit, Zymo Research) were reversed transcribed using MMLV reverse transcription (ThermoFisher Scientific), qPCR analyses were performed using SensiFastSybr Lo-Rox Mix (Bioline) in Applied Biosystems QuantStudio thermal cycler using the relative standard curve method (Applied Biosystems User Bulletin #2). All conditions were previously described [1].

Oligonucleotides for export and total RNA levels:

Myc forward: 5' CTTCTCTGAAAGGCTCTCCTTG 3'

Myc reverse: 5’ GTCGAGGTCATAGTTCCTGTTG 3’

MCL1 forward: 5’ GGCAGTCGCTGGAGATTAT3’

MCL1 reverse: 5’ CAACCCGTCGTAAGGTCTC3’

β-Actin forward: 5’ GCATGGAGTCCTGTGGCATCCACG3’

β-Actin reverse: 5’ GGTGTAACGCAACTAAGTCATAG3’

GAPDH forward: 5’ GAAGGTGAAGGTCGGAGTC3’

GAPDH reverse: 5’ GAAGATGGTGATGGGATTTC3’

U6 forward: 5’ CGCTTCGGCAGCACATATAC 3’

U6 reverse: 5’ AAAATATGGAACGCTTCACGA 3’

tRNAMet forward: 5’ AGCAGAGTGGCGCAGCGG 3’

tRNAMet reverse: 5’ GATCCATCGACCTCTGGGTTA 3’

**7. Flow cytometry**

Single cell suspensions were stained with the following phycoerythrin (PE), PE-CY7, fluorescein isothiocyanate, allophycocyanin (APC), APC-Cy7, or Pacific blue conjugated antibody clones(BioLegend, San Diego,CA): Mac-1(M1/70), Gr1 (RB6-8C5), CD4 (GK1.5), CD8 (53-6.7), B220 (RA3-6B2), Sca1 (D7), c-Kit (2B8), CD45.1 (A20) and CD45.2(104). Cells were analyzed with an LSRII flow cytometer (BD Biosciences) or FACSAria (BD Biosciences) and sorted using a FACSAria.

Diva software (BD) and FlowJo (Tree Star) were used for data acquisition and analysis, respectively.

### 8. Serial replating assay

The serial replating assay was performed as described [4] with the following deviations. BMCs for retroviral transduction were harvested from 5-FU treated B6.SJL-Ptprca Pepcb/BoyJ mice. Transduction was performed using Retronectin (Takara Bio Inc.) coated 35mm dishes according to the manufacturer. Replating was carried out for 4 consecutive rounds until the proliferative capacity of controls was largely exhausted.

### 9. Plasmids

The MSCV-IRES-GFP (MIG)-flag-HoxA9 vector was a kind gift from Scott Armstrong5. pMSCV-IRES-YFP (MIY) (Addgene plasmid # 52108) was originally described in 6. The Meis1 open reading frame was cloned from pMSCV-puro-Meis1 (a gift from Scott Armstrong). HoxA9Y11AL16A (HoxA9AA) was created by successive application of the QuikChange site-directed mutagenesis kit (Agilent, La Jolla, CA) to the MIG-flag-HoxA9 vector using the following oligos (mutated sites in bold):

HoxA9Y11Amutfw: CTGGGCAACTAC**gc**TGTGGACTCCTTC

HoxA9Y11Amutrev: GAAGGAGTCCACA**gc**GTAGTTGCCCAG

HoxA9L16Amutfw:GTGGACTCCTTC**gcc**CTGGGCGCCGAC

HoxA9L16Amutrev:GTCGGCGCCCAG**ggc**GAAGGAGTCCAC

The N51S non-DNA binding mutant of HoxA9 was described earlier [7] and was introduced into the MIG-flagHoxA9 vector using the QuikChange Kit with the following oligos (mutated site in bold):

HoxA9_N51Sfw:

GCAGGTCAAGATCTGGTTCCAGA**g**CCGCAGGATGAAAATGAAG

HoxA9_N51Srev:

CTTCATTTTCATCCTGCGG**c**TCTGGAACCAGATCTTGACCTGC

**10. RNA preparation from mouse BMCs**

BMCs from each of 3 Hoxa9wt/Meis1and Hoxa9AA/Meis1 were harvested with ≥ 97% GFP+ cells in the leukemic bone marrow. Total RNA was extracted with TriReagent (Sigma-Aldrich, St. Louis, MO) and purified with RNeasy Micro Plus columns in conjunction with gDNA Eliminator columns (Qiagen, Hilden Germany) to remove residual genomic DNA. RNA samples were analyzed using Gene Chip Mouse Gene 1.0 Assay Kit.

### 11. Electrophoretic mobility shift assay (EMSA)

Annealed DNA oligonucleotides containing a HoxA9 consensus binding motif (15 pmol, sense: 5‘-ctgcgatgatttacgaccgc-3‘, antisense: 5‘-gcggtcgtaaatcatcgcag-3‘, described in [8] were end-labelled with[γ-32P] ATP (Perkin Elmer) and T4 polynucleotide kinase (New England Biolabs). Reactions were setup essentially as described earlier [9]with the exception that nuclear lysates from HoxA9, HoxA9AA, and HoxA9N51S transfected HEK293T were used in the binding reactions. Nuclear extracts were prepared using the NE-PER Nuclear and Cytoplasmic Extraction Reagents (Pierce, Thermo Fisher Scientific, Waltham, MA). Briefly,10µg of nuclear extract were incubated with labelled DNA oligonucleotide in binding buffer (5mM Tris, pH 7.4, 5mM MgCl2, 1mM dithiothreitol (DTT), 3% (v/v) glycerol, 100mM NaCl). Supershift reactions were performed by adding 2ug of anti-flag M2 antibody. Cold competition reactions were carried out by adding 100-fold molar excess of non-radioactive labelled dsDNA probe.

### Supplemental Figures

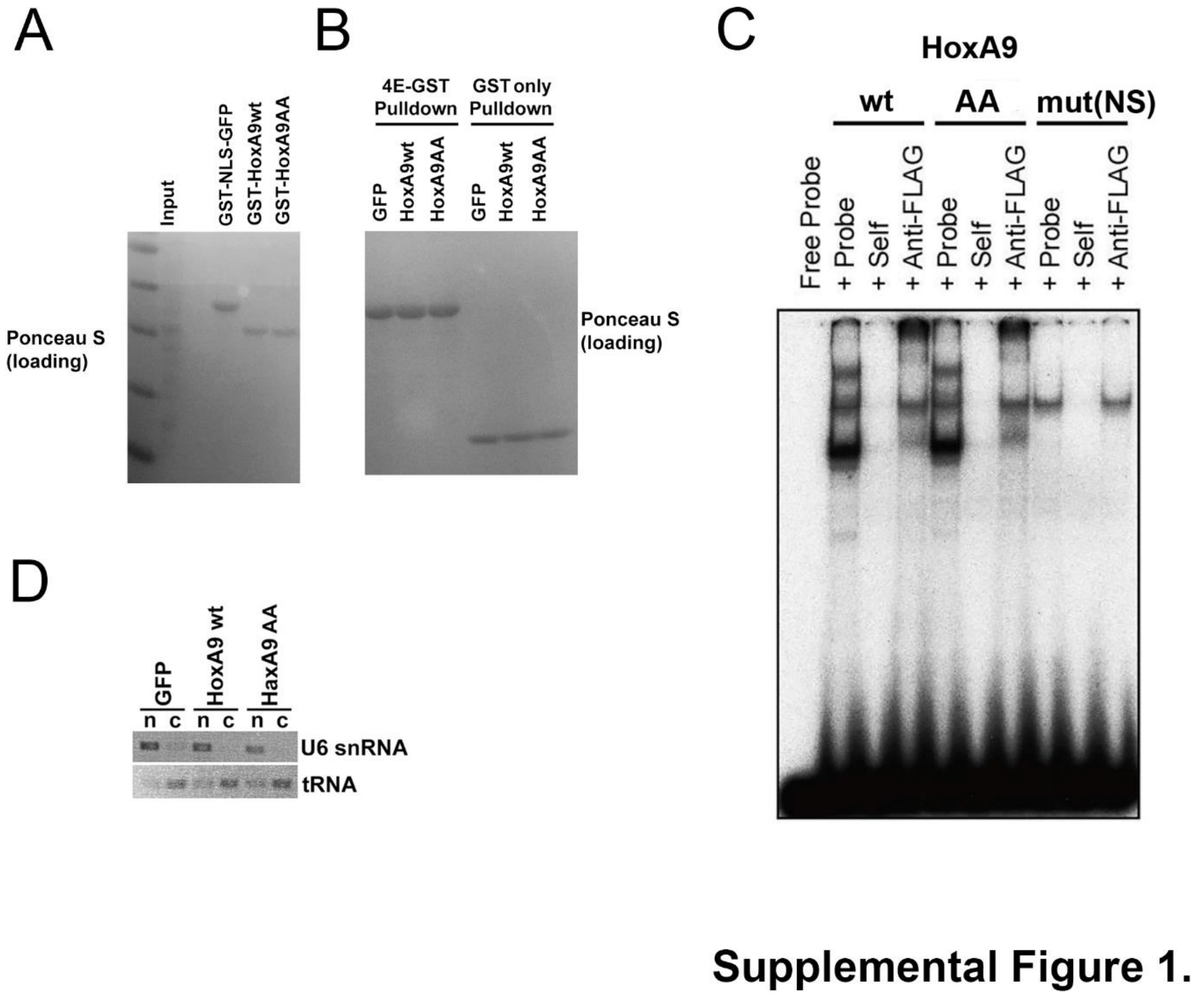


**Figure S1. The impacts of HoxA9 mutation on eIF4E and DNA in K562 cells.**

(A-B) Loading controls for GST pulldowns. (A) Ponceau S of immunoblot from Figure 1B. (B) Ponceau S of immunoblot for Figure 1C. (C) Electrophoretic Mobility Shift Assay (EMSA) for-HoxA9wt, HoxA9AA, and HoxA9 DNA binding mutant (NS). (D) representative semi-qRT-PCR for U6 snRNA and $tRNA_{Met}$ as controls for the nuclear (n) and cytoplasmic (c) fractions respectively, corresponding to mRNA export panel in Figure 1D.

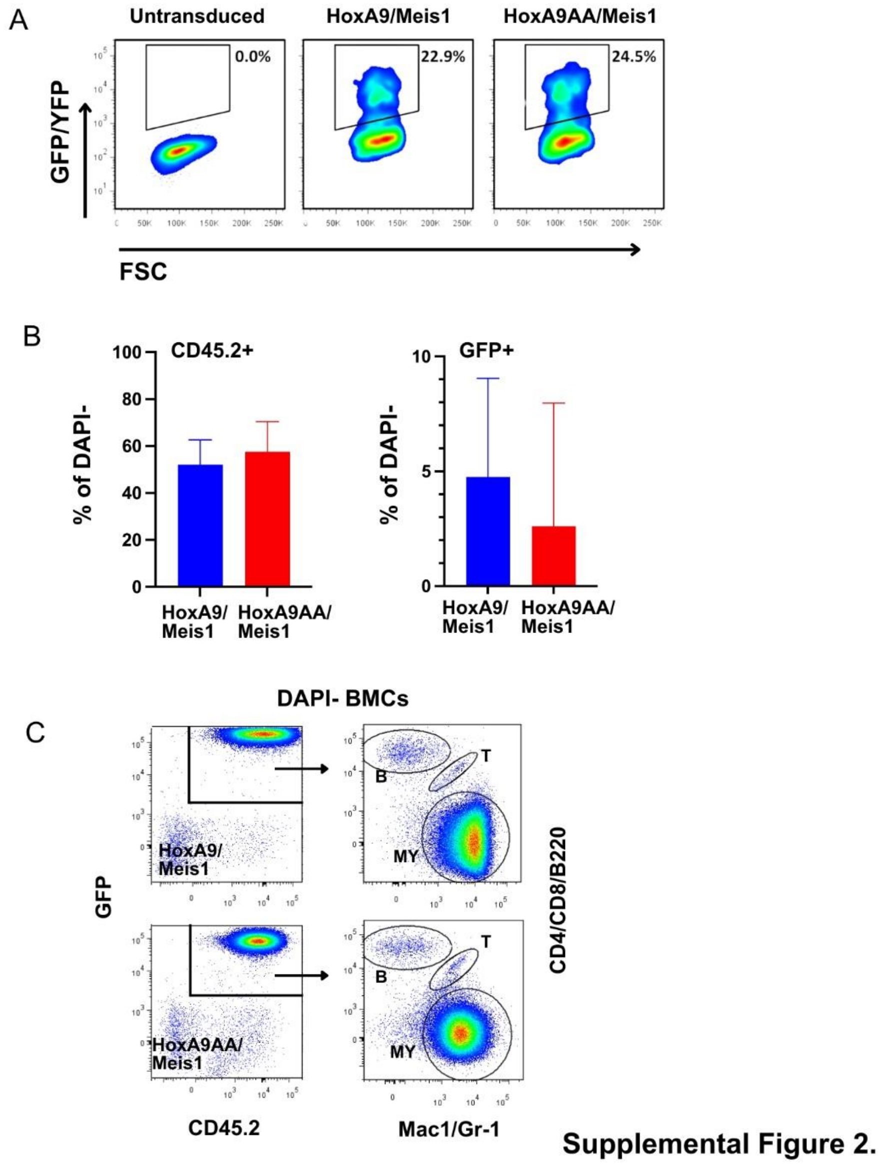


**Figure S2. Flow cytometric analysis of transplanted HoxA9wt/Meis1 and HoxA9AA/Meis1 mice.** (A) Efficiency of HoxA9wt/Meis1 and HoxA9AA/Meis1 transduction of KSL cells at the time of transplantation. (B) Engraftment efficiency 4 weeks post-transplantation measured by flow cytometric analysis of peripheral blood. (C) Flow cytometric characteristic bone marrow cells of diseased mice after transplantation. MY indicates myeloid cells.
